# Serial cryoFIB/SEM reveals profound cytoarchitectural disruptions caused by a pathogenic mutation in Leigh syndrome patient cells

**DOI:** 10.1101/2020.05.28.121111

**Authors:** Yanan Zhu, Dapeng Sun, Andreas Schertel, Jiying Ning, Xiaofeng Fu, Pam Pam Gwo, Alan M. Watson, Zachary Freyberg, Peijun Zhang

**Affiliations:** Division of Structural Biology, Wellcome Trust Centre for Human Genetics, University of Oxford, Oxford, OX3 7BN, UK; Department of Structural Biology, University of Pittsburgh School of Medicine, Pittsburgh, PA 15260, USA; Carl Zeiss Microscopy GmbH, Zeiss Customer Center Europe, Carl-Zeiss-Straße 22, D-73447 Oberkochen, Germany; Department of Psychiatry, University of Pittsburgh, Pittsburgh, PA 15213, USA; Department of Cell Biology, University of Pittsburgh, Pittsburgh, PA 15213, USA; Electron Bio-Imaging Centre, Diamond Light Source, Harwell Science and Innovation Campus, Didcot OX11 0DE, UK

**Author notes:** These authors contributed equally (Yanan Zhu, Dapeng Sun).

## Abstract

The advancement of serial cryo-FIB/SEM offers a new opportunity to study large volumes of near-native, fully hydrated frozen cells and tissues at voxel sizes of 10 nm and below. We explored this capability for pathologic characterization of vitrified human patient cells. We demonstrate profound disruption of subcellular architecture in primary fibroblasts from a Leigh syndrome patient harboring a disease-causing mutation in USMG5 protein responsible for impaired mitochondrial energy production.

## Introduction

The primary role of mitochondria is to generate energy in cells through mitochondrial oxidative phosphorylation (OXPHOS)[1]. OXPHOS deficiency leads to mitochondrial diseases including Leigh syndrome (LS), a progressive neurological disorder and the most common mitochondrial disease in children[2]. LS is genetically heterogeneous with more than 90 nuclear or mitochondrial genes implicated in its pathogenesis. Virtually all of these genes encode the mitochondrial respiratory complex machinery required for energy generation through OXPHOS[3] including those regulating the structure and assembly of complex V (ATP synthase). Recent studies identified a novel pathogenic mutation, (c.87 + 1G>C), in the *USMG5* gene that results in autosomal recessive LS[3]. As previously described, the mutation abolishes the canonical GT splice site donor of Exon 4 of *USMG5* and produces aberrant transcripts that are degraded via nonsense-mediated decay with >90% loss of USMG5 expression[3]. USMG5, also known as DAPIT (diabetes-associated protein in insulin-sensitive tissues), is a constituent of complex V required for its dimerization. Complex V ordinarily exists as a dimeric supercomplex required to shape the mitochondrial cristae, enabling efficient flow of the protons needed to fuel ATP synthesis. Significantly, recent cryo-electron tomography (cryoET) studies of LS patient cells harboring this *USMG5* gene mutation revealed significant disturbances in the ultrastructure of mitochondrial crista, with profound overall loss of cristae and dilatation of the remaining few cristae[4]. The effect of the *USMG5* mutation on the whole cell and subcellular architecture, however, has not been investigated.

CryoET, with subtomogram averaging, has emerged as a powerful method for visualizing heterogeneous structures and *in situ* specimens at subnanometer resolutions[5, 6]. However, a major limitation of cryoET’s utility is its reliance on very thin samples (<300nm), such as thin regions of the cell periphery or cell lamella by cryo-FIB thinning, due to limited penetrance of the electron beam in thicker regions of cells [7, 8]. On the other hand, serial FIB/SEM has been rapidly adopted as a technique for generating large 3D volumes, including multiple cells and tissue constituents, of fixed (cryo or chemically), dehydrated, resin-embedded and stained samples[9, 10]. Its application to vitreous biological samples, namely serial cryoFIB/SEM, involves the challenges associated with cryo-electron microscopy and is an emerging, promising method for studying plunge-frozen and high pressure frozen cells and tissues[11–18]. The potential of combining large scale cryo-volume imaging using serial cryoFIB/SEM, followed by cryoFIB lamella preparation of the specific region of interest identified through serial cryoFIB/SEM, with cryoET imaging of the targeted lamella at a high resolution on the exact same specimen, is especially exciting.

Here, we applied state-of-the-art serial cryoFIB/SEM to plunge-frozen primary fibroblast cells from a healthy individual and from an LS patient carrying the homozygous mutation in the *USMG5* gene previously shown to impair complex V dimerization, cristae structure and ATP synthesis[4]. We explored the application of this emerging new capability for clinical phenotyping of pathological cells in close-to-native state and demonstrate a profound disruption of subcellular structures in patient cells. Compared to conventional serial FIB/SEM of stained and resin-embedded samples, serial cryoFIB/SEM offers a much faster (without dehydration and embedding steps) and near-native technique for phenotypic characterization of whole-cells or tissue, which could be exceedingly useful in clinical settings.

## Results and discussion

To investigate the phenotypic impact of a specific *USMG5* gene mutation (c.87 + 1G>C) on cellular and subcellular structures in a near-native state, we used serial cryoFIB sectioning and cryoSEM imaging of frozen-hydrated primary fibroblast cells isolated from an LS patient and from a healthy individual (Table S1). Using a lateral pixel spacing of 10.5 nm for SEM imaging and a FIB slice thickness of 21 nm, an entire patient fibroblast cell was sliced through and 2018 slices were imaged at 4096 × 3072 pixel in about ~17.5 hours, resulting in a total volume of 58789 μm^3^ (Movie S1). For the control fibroblast cell, similar voxel size was used, but a reduced raster of 3072 x 1150 pixel was used for imaging in total 575 slices since the cell width was much larger than its height. A total volume of 4062 μm^3^ was obtained within ~5.5 hours for the control cell (Movie S2). Figure 1 shows SEM images of representative slices from both patient and control cells. We noted a contrast imbalance between the lower part and the upper part of the cross-sectioned patient cell (Figure 1j). Since the SEM image contrast is related to local surface potentials, the median potential (or threshold) can be different depending on the environment and local charge dissipation, such as the top cell region being close to the cold deposited Platinum precursor layer and bottom part being in the vicinity of the support film, grid bars and top surface of the cryo-holder. In both cells, subcellular structures are clearly visible, especially membrane-enclosed sub-cellular compartments (Figure 1a&j) including the nucleus. Notably, serial cryoFIB/SEM provides unambiguous visualization of nuclear pores (Figure 1j-l, arrowheads). In the control cell, individual cellular organelles can be identified based on their distinct morphologies, including the cell membrane, a phagosome (Figure 1b-d, PG, in three consecutive slices), endoplasmic reticulum (ER), multivesicular bodies (Figure 1e-f, MV), Golgi (Figure 1g, *), mitochondria (Figure 1g-i, white arrows), vacuole-like membranous structures (Figure 1i, V) and the cell nucleus. The LS patient fibroblast cell, however, shows substantial cytoarchitectural derangements, with the interior of the cell largely occupied by vacuolated structures of indeterminate origin. Of the residual identifiable structures including mitochondria and Golgi, organelles are significantly decreased in volume and displayed gross morphological abnormalities. For example, the Golgi apparatus lacks extended membrane stacks (Figure 1j&m). More remarkably, compared to the complex shape and network of mitochondria in the control cell (Figure 1a, g, h, Movie S2), nearly all patient mitochondria are roughly round with minimal cristae (Figure 1j, l, n, Movie S1), consistent with our previous cryoET analyses[4]. This suggests that, consistent with the earlier biochemical characterization of these primary fibroblasts[3], the architecture responsible for energy metabolism in the patient cells is compromised. The patient cells also grow substantially slower than the control cells.

**Figure 1.**
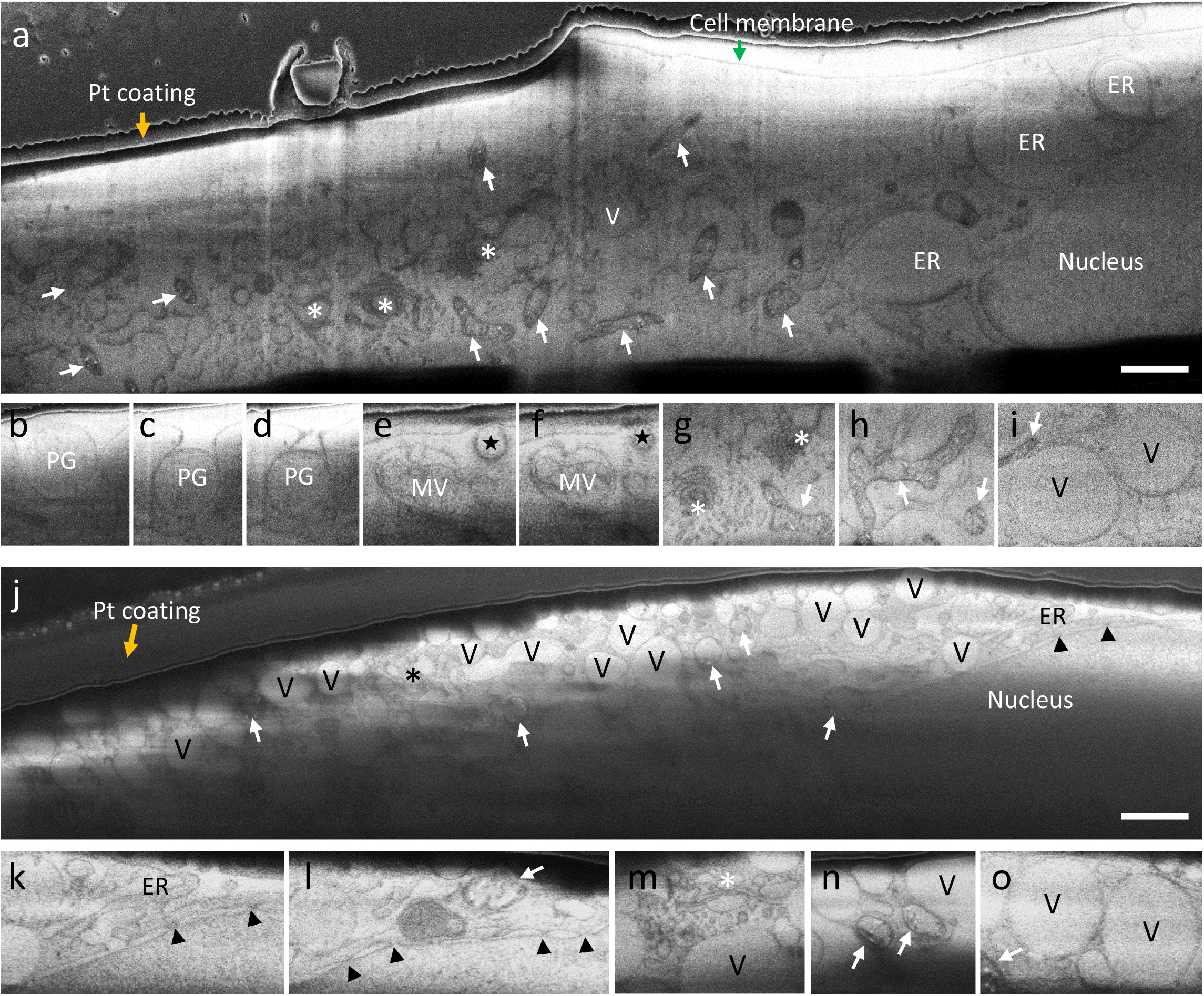
Serial cryoFIB/SEM of frozen-hydrated primary cells from control and patient fibroblasts. a) A representative cryoSEM image from a stack of 575 serial micrographs recorded from a control fibroblast cell cultured on an EM grid. Arrows, mitochondria; asterisks, Golgi; stars, endosome; PG, phagosome; MV, multivesicular body; V, vacuoles; arrowheads, nuclear pore; orange arrows, platinum GIS coating; green arrow, cell membrane; ER, endoplasmic reticulum. b-i) An image gallery of subcellular structures and organelles observed in the control cell, including three consecutive slices of a phagosome entering the cell (b-d), two consecutive slices of an endosome (e-f, star), a multivesicular body (e-f, MV), Golgi complexes (g, asterisks), tubular shaped mitochondria (g-h, arrow), and vacuoles (i, v). j) A representative cryoSEM image from a stack of 2018 serial micrographs recorded from a LS patient fibroblast cell cultured on an EM grid. k-o) An image gallery of subcellular structures and organelles observed in the patient cell, showing endoplasmic reticulum (k, ER), nuclear pores (k-l, arrowheads), Golgi complex (m, asterisk), mitochondria (l&n, arrows) and vacuoles (n-o, V). Scale bars, 1 μm.

For further analysis, we have performed 3D reconstruction and segmentation of the serial cryoFIB/SEM data for both patient and control cells (Figure 2a&b, Movies S1&S2). In the volume rendering of the control cell, an extended network of oblate tubular-shaped mitochondria is evident (Figure 2a, green, Movie S3), whereas the patient cell shows mitochondria that are not continuous and mostly round or oval-shaped individuals (Figure 2b, green, Movie S4). Some individual mitochondria appear in close association with one another in both control and patient cells (Figure S1, Movie S7 and S8). The size of vacuoles (Figure 2b, yellow) within the patient cell are also remarkably larger than those of the control cell and are more densely packed in the patient cell (Figure 1a&j, Figure 2a-b). To analyze the structural details of organelles, a small region of the cell was cropped and segmented semi-automatically, as shown in Figure 2c&d (Movies S5&S6). We can appreciate the drastic differences in the morphology of mitochondria and the shape and distribution of cristae between patient and control cells. Cristae structure is severely disturbed in the patient cell, appearing sparse in number and short, as previously observed by cryoET of limited regions of the cell periphery[4]. The dramatic impact of Complex V’s failure to dimerize due to a specific *USMG5* gene mutation on overall cellular architecture and organelle structures in LS patient cells is now more fully appreciated in the greater context of the whole cell though *in situ* large volume imaging.

**Figure 2.**
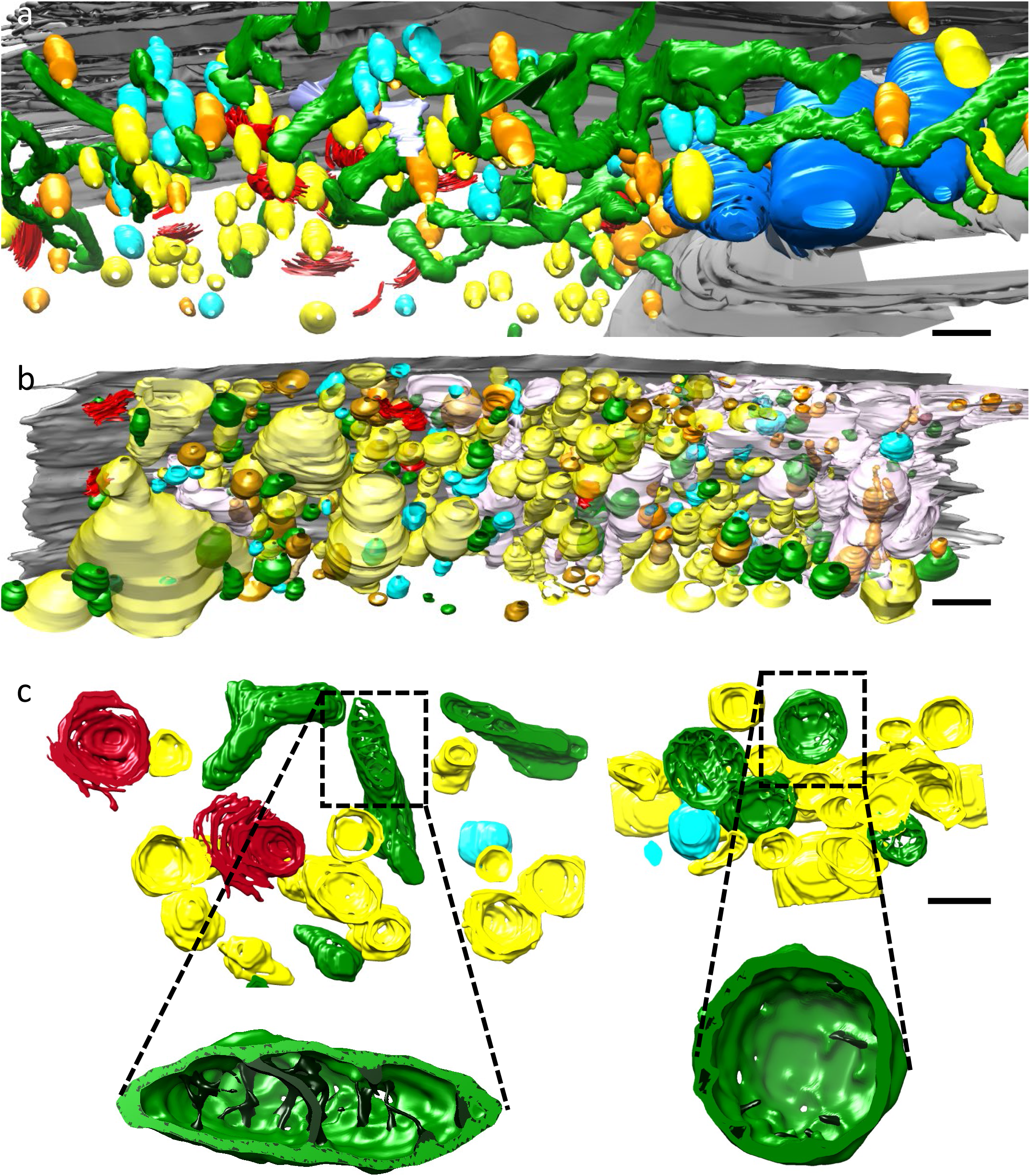
3D reconstruction and segmentation of control and patient cells. a-b) Surface rendering of segmented volumes of control (a) and patient (b) fibroblast cells. Green, mitochondria; red, Golgi, yellow, vacuoles; orange, dense vesicles; cyan, partially dense vesicles, Blue, ER. c) Segmentation of a small representative volume from control (left) and patient (right) fibroblasts. Inserts are enlarged views of a single mitochondrion in control (left) and patient (right) cells. Cristae are shown in dark green. Scale bars, 1 μm in a&b, 500 nm in c.

The majority of imaging studies on LS disease have been understandably focused on mitochondria. Nevertheless, our work demonstrates dramatic phenotypic changes to LS patient cells that extend beyond mitochondria to alter most, if not all, organelles within the cell. Using emerging serial cryoFIB/SEM technology, we captured and visualized an entire frozen-hydrated mammalian cell in 3D for the first time. More importantly, we applied this capability to studies of human cellular disease processes. This revealed a profound disruption of cellular and subcellular structures in a primary LS patient fibroblast. Such whole cell volume phenotypic characterization of cells and tissues *in situ*, at the near-native state, offers the opportunity to improve our understanding of diseases beyond LS and potentially provides new means for clinical use, from diagnosis to treatment.

### Methods

The methods section is linked to the online version of the paper at https://www.nature.com/nmeth/

## Data availability

The raw serial cryoFIB/SEM data from both control and patient cells will be deposited in the EMPIAR database.

## Acknowledgements

We thank Dr. Teresa Brosenitsch for critical reading of the manuscript. We are grateful to Dr. Michio Hirano for providing the control and patient cells. We also thank Drs. James Gilchrist for technical support, and Drs. Tao Ni, Min Xu, Julika Radecke and Andrew Howe for helpful discussions about segmentation. We thank Dr. Saskia Mimietz-Oeckler and Andreas Halladay, Leica Microsystems for technical support. This work was supported by the Department of Defense grants PR141292 (Z.F.) and PR192466 (Z.F.), the National Institutes of Health P50 grant AI150481 (P.Z.), the UK Wellcome Trust Investigator Award 206422/Z/17/Z (P.Z.), and the UK Biotechnology and Biological Sciences Research Council grant BB/S003339/1 (P.Z.).

## Author contributions

P.Z. conceived and designed the experiments. J.N. cultured cells and X.F. prepared cryo-specimens. A.S. performed serial cryoFIB/SEM. Y.Z., P.P.G., A.M.W. and D.S. performed segmentation and, with P.Z., analyzed data. Y.Z. and P.Z. wrote the paper with support from other authors.

## Online Methods

### Sample preparation

Skin fibroblasts from the patient, expressing a homozygous *USMG5* mutation (C.87+1G>C, 1 base pair after the end of Exon 3), and from a healthy human subject (provided by Dr. Michio Hirano, Columbia University) were cultured in Dulbecco’s minimal essential media (DMEM) (Thermo Fisher, Waltham, MA, USA) supplemented with 15% fetal bovine serum (FBS) (Sigma-Aldrich, St. Louis, MO, USA), 1% vitamin solution and 1% antibiotic-antimycotic (Thermo Fisher) as described earlier[4]. All experiments were conducted on cells cultured for <15 passages. Cells were plated onto gold R2/2 Quantifoil finder EM grids (Quantifoil Micro Tools GmbH, Jena, Germany) at density of 0.5-1 × 10^5^ cells/ml (total 2 ml culture) in glass-bottom culture dishes (MatTek Corporation, Ashland, MA). The gold EM grids were coated with 50 μg/ml fibronectin (Sigma-Aldrich) and sterilized under UV light for 2 hours before use. For the control cells, after 48 hours culture, the grids were blotted with a filter paper and plunge-frozen into liquid ethane for rapid vitrification using an FEI Vitrobot (FEI, Hillsboro, OR) at ~100% humidity. Patient cells grew slowly and were cultured for 5 days before plunge-freezing.

### Serial cryoFIB/SEM

For patient cells, the plunge-frozen EM grid was mounted in a Leica Vacuum Cryo Manipulation (VCM) preparation box (Leica Microsystems GmbH, Vienna, Austria) on a Leica cryo-holder for freeze-fracturing under cryogenic conditions. The TEM grid was held down on the flat top surface by using the clamp. The sample holder was transferred into a Leica ACE 600 cryo-sputter coater using a Leica VCT500 sample shuttle (Leica Microsystems, Vienna, Austria). At −154 °C, the sample was sputter-coated with a 4 nm thick tungsten layer. The samples were then transferred into a ZEISS Crossbeam 550 FIB-SEM (Carl Zeiss Microscopy GmbH, Oberkochen, Germany). The cryo-stage temperature was maintained at −155 °C and the system vacuum was 1.6×10^−6^ mbar. For areas containing cells, a cold deposition of platinum precursor material was achieved by opening the gas valve for 45 seconds. For cold deposition, the gas reservoir temperature was 28 °C, and the distance between the gas capillary and the sample was about 3 mm.

First, a viewing channel for SEM imaging was milled using a FIB-milling probe current of 7 nA. The resulting cross-section was polished with a FIB probe current of 3 nA. For serial sectioning and imaging, a FIB probe current of 700 pA was used, and the FIB slice thickness was 21 nm. SEM images using InLens SE detection with a SEM probe current of 35 pA, a SEM acceleration potential of 2.3 kV and a dwell time of 100 ns were recorded. The lateral pixel spacing for SEM imaging was 10.5 nm and the image size was 4096 × 3072 pixels. Line Average with a line average count N = 61 was used for noise reduction.

The control cell sample was mounted on a pre-tilted Leica sample holder for on-grid-thinning (Leica Microsystems GmbH, Vienna, Austria). After sputter-coating and transfer into the Crossbeam 550 FIB/SEM, a cold deposition of platinum precursor was done following the same procedure as above. The Crossbeam 550 system pressure was 8×10^−7^ mbar. A FIB probe current of 700 pA and a FIB slice thickness of 20 nm were used for serial sectioning and imaging. The lateral pixel spacing for SEM imaging was 10 nm and the imaging box was reduced to 3072 × 1150 pixel since the cell width is much larger than its height. For SEM imaging, InLens SE detection, a SEM probe current of 59 pA, a SEM acceleration potential of 1.9 kV and a dwell time of 200 ns were used. We employed a line average count N = 19 was used for noise reduction.

### Local Reconstruction and segmentation of 3D subvolumes

The raw SEM images were first aligned and a 3D volume generated using IMOD[19]. Subvolumes with representative features were cropped out from each aligned image stack (300×200 pixel × 40 slices or ~3.2 × 2.1 × 0.84 μm for patient cell and 500×400 pixel×40 slices or ~5.0 × 4.0 × 0.8 μm for control cell). The alignment between images was refined using an in-house Matlab script based on the Imregister function (Supplementary material). To make objects smooth, 19 additional images were generated, with a linear interpolation, and inserted between two successive image slices with a home-made Matlab script (Supplementary material). Using PixelAnnotationTool available online (https://github.com/abreheret/PixelAnnotationTool/releases), specific organelles, such as mitochondria, Golgi etc., were labeled manually from the cropped images as masks, which were then used to extract the image data from the corresponding region separately. These segmented image volumes were displayed in Chimera[20].

### Global image alignment and 3D modelling

The region containing the cell content was masked using Chimera software[20] with a boundary contour that was generated in 3dmod to just include the cell. The masked image slices (237 slices from the control cell and 218 slices from the patient cell) were first aligned with the Tiltxcorr[19] program in IMOD using cross-correlation to determine the X and Y translations between successive image slices. The coarse-aligned image stacks were registered further using a SIFT-based algorithm[21] adapted to run on TrakEM2 plugin for FIJI[22]. The post-registration images were exported into 3dmod[19] for further manual segmentation. Chimera was used for display of segmented 3D models.

---

**Figure S1.**
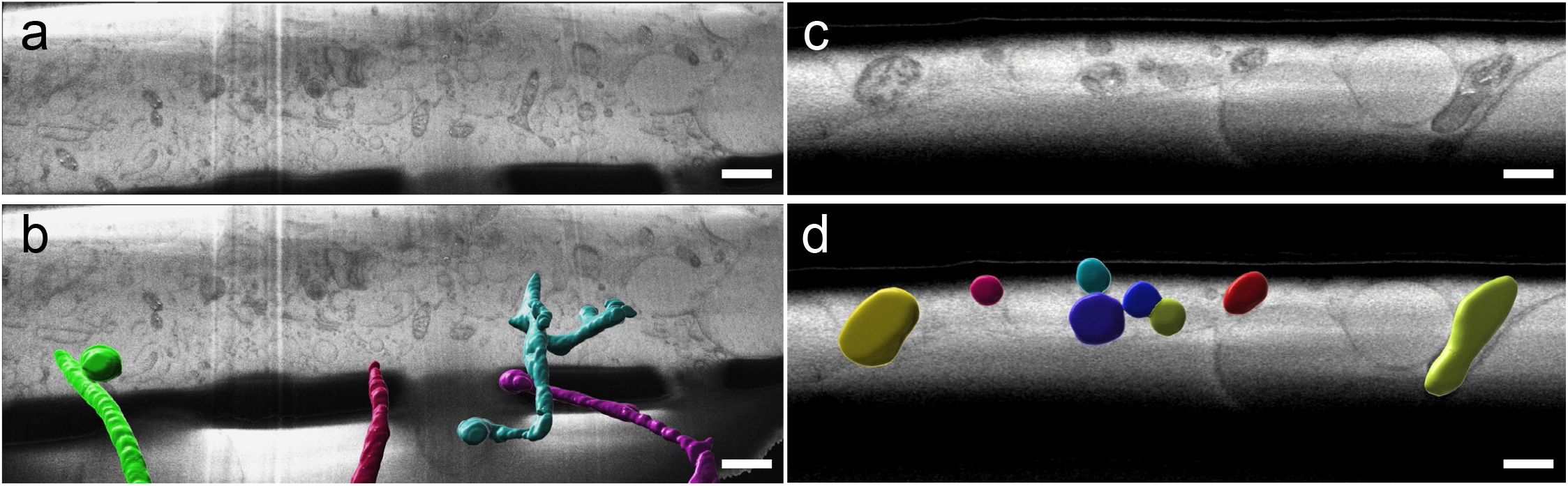
Close association of mitochondria in control (a-b) and patient (c-d) cells. Mitochondria were manually segmented from control (a) and patient (c) datasets using NIS Elements (Nikon). The segmented images were overlaid with the RAW data in Imaris 9.5 (Bitplane) where the segmented areas were transformed into objects using surfacing tools (b and d, also see Movie S7 and S8). Independent objects are signified by a unique color. Scale bars, 1 μm in a & b, 0.3 μm in c & d.

**Table S1:** Summary of serial cryoFIB/SEM parameters.

|  | **Pixel size (nm)** | **Image size (pixel)** | **No. Slices** | **Volume ($\mu\text{m}^3$)** | **Time (h:m)** |
| --- | --- | --- | --- | --- | --- |
| **Patient** | 10.5 x 10.5 x 21 | 4096 × 3072 | 2018 | 50,784 | 17:20 |
| **Control** | 10 x 10 x 20 | 3072 × 1150 | 575 | 4,062 | 5:40 |

## Movie legends

Movie S1. Aligned raw image slices of patient fibroblast, overlaid with segmentation.

Movie S2. Aligned raw image slices of control fibroblast, overlaid with segmentation.

Movie S3. Segmented representation of control fibroblast.

Movie S4. Segmented representation of patient fibroblast

Movie S5. A subregion of aligned raw image slices of control fibroblast, overlaid with segmentation.

Movie S6. A subregion of aligned raw image slices of patient fibroblast, overlaid with segmentation.

Movie S7. Segmented region of mitochondria from control fibroblast demonstrating close contacts.

Movie S8. Segmented region of mitochondria from patient fibroblast demonstrating close contacts.

